# Beer ethanol and iso-α-acid level affect microbial community establishment and beer chemistry throughout wood maturation of beer

**DOI:** 10.1101/2022.03.07.483260

**Authors:** Sofie Bossaert, Tin Kocijan, Valérie Winne, Johanna Schlich, Beatriz Herrera-Malaver, Kevin J. Verstrepen, Filip Van Opstaele, Gert De Rouck, Sam Crauwels, Bart Lievens

## Abstract

Sour beers produced by barrel-aging of conventionally fermented beers are becoming increasingly popular. However, as the intricate interactions between the wood, the microbes and the beer are still unclear, wood maturation often leads to inconsistent end products with undesired sensory properties. Previous research on industrial barrel-aging of beer suggests that beer parameters like the ethanol content and bitterness play an important role in the microbial community composition and beer chemistry, but their exact impact still remains to be investigated. In this study, an experimentally tractable lab-scale system based on an *in-vitro* community of four key bacteria (*Acetobacter malorum*, *Gluconobacter oxydans*, *Lactobacillus brevis* and *Pediococcus damnosus*) and four key yeasts (*Brettanomyces bruxellensis*, *Candida friedrichii*, *Pichia membranifaciens* and *Saccharomyces cerevisiae*) that are consistently associated with barrel-aging of beer, was used to test the hypotheses that beer ethanol and bitterness impact microbial community composition and beer chemistry. Experiments were performed using different levels of ethanol (5.2 v/v%, 8 v/v% and 11 v/v%) and bitterness (13 ppm, 35 ppm and 170 ppm iso-α-acids), and beers were matured for 60 days. Samples were taken after 0, 10, 20, 30 and 60 days to monitor population densities and beer chemistry. Results revealed that all treatments and the maturation time significantly affected the microbial community composition and beer chemistry. More specifically, the ethanol treatments obstructed growth of *L. brevis* and *G. oxydans* and delayed fungal growth. The iso-α-acid treatments hindered growth of *L. brevis* and stimulated growth of *P. membranifaciens*, while the other strains remained unaffected. Beer chemistry was found to be affected by higher ethanol levels, which led to an increased extraction of wood-derived compounds. Furthermore, the distinct microbial communities also induced changes in the chemical composition of the beer samples, leading to concentration differences in beer- and wood-derived compounds like 4-ethyl guaiacol, 4-ethyl phenol, cis-oak lactone, vanillin, furfural and 5-methyl furfural. Altogether, our results indicate that wood-aging of beer is affected by biotic and abiotic parameters, influencing the quality of the final product. Additionally, this work provides a new, cost-effective approach to study the production of barrel-aged beers based on a simplified microbial community model.

## 1 Introduction

Sour beers produced by wood-aging of conventionally fermented beers are receiving an increased interest in the (craft) beer community due to their noteworthy balance between sourness, aroma and flavor complexity. Typically, following conventional fermentation these beers are aged for a long period of time (often up to several months, a year, or more) in wooden barrels leading to beers with a unique sour flavor palette (Bossaert *et al.*, 2019). Besides the extraction of wood-derived compounds into the beer, wood-aged sours owe their layered flavor profile to the metabolic activity of a variety of ‘wild’ microorganisms that subside within the pores of wooden barrels or that originate from the brewery environment (Bossaert *et al.*, 2021a; 2021b; De Roos *et al.*, 2019). These microorganisms generally include conventional yeasts belonging to the genus of *Saccharomyces* and various ‘wild’ fungi belonging to the genera *Brettanomyces*, *Candida*, *Debaryomyces*, *Pichia*, and *Wickerhamomyces* (Bossaert *et al.*, 2021a; 2021b; Osburn *et al.*, 2018), in addition to a number of lactic acid bacteria and acetic acid bacteria (De Roos and De Vuyst, 2019). Whereas lactic acid and acetic acid are the main contributors to the beer’s sourness (Bossaert *et* al., 2019; De Roos and De Vuyst, 2019), other common microbial metabolites found in wood-aged sour beers include 4-vinyl guaiacol, 4-ethyl guaiacol and 4-ethyl phenol that impart a spicy and barnyard aroma to the beer (De Roos and De Vuyst, 2019), while higher alcohols and esters are mainly responsible for the floral and fruity flavor notes (Dzialo *et al.*, 2017). In general, the process of wood-aging and the final beer characteristics are determined by many factors like wood species, geographic origin of the wood, toasting degree of the wood, wood history, barrel dimensions, characteristics of the beer aging inside, characteristics of other beverages that were previously matured inside the barrel, ambient conditions (e.g. temperature and humidity), brewing environment, and duration of the maturation (Bossaert *et al.*, 2021a; 2021b; Fan *et al.*, 2006; Sterckx *et al.*, 2012a; 2012b). However, so far only little is known about the exact contribution of each of these parameters to the final beer, particularly because their effects have not yet been studied systematically. As a result, wood-aging of beer largely remains a black-box process to date, which often generates undesirable sensory characteristics of the end product or inconsistency in quality, leading to economic losses. Therefore, there is a clear need for a better understanding of the wood-aging process, including the interactions that take place between the environment, the wood, the microorganisms and the maturing beer.

Previous research performed on an industrial scale suggests that beer parameters like ethanol content and beer bitterness play an important role in the composition of the microbial community and the beer chemistry of sour beers produced via wood-aging (Bossaert *et al.*, 2021a). More specifically, Bossaert *et al.* (2021a) indicated that ethanol level affected the fungal community composition and the extraction of wood compounds, while beer bitterness restrained the bacterial community composition. However, to assess the true effects of these parameters, a more fundamental, in-depth study under controlled conditions is required in which several levels of these parameters are assessed, while all other factors remain constant. Nevertheless, performing such studies in a statistical meaningful manner in industrial conditions is challenging as large quantities of wooden barrels need to be purchased and filled with beer. Furthermore, the barrels should be stored under the same experimental conditions and sampled over a long period of time. One way to overcome these challenges is to use a small scale, experimentally tractable model system based on a number of key microorganisms that mimics industrial wood-aging and that can be manipulated to assess the influence of the parameters of interest. In recent years, similar model systems have been used to get a better understanding of the assembly and function of microbial communities (Cosetta *et al.*, 2020; Jia *et al.*, 2020; Wolfe *et al.*, 2014; Wolfe and Dutton, 2015; Wolfe, 2018). For example, using an *in-vitro* microbial community of 11 key microbes occurring on the surface of cheese during aging (also known as the ‘cheese rind’), Wolfe *et al.* (2014) unambiguously showed that abiotic parameters like moisture strongly affect microbial community formation. Previous work has provided a detailed view on the dynamics of the microbial community composition in the wood-aging of beer. Beers were found to undergo a shift from a diverse microbial community to a number of core microorganisms, including lactic acid bacteria like *Lactobacillus* and *Pediococcus*, acetic acid bacteria like *Acetobacter*, and wild yeasts like *Brettanomyces* and *Pichia* (Bossaert *et al.*, 2021a; 2021b). Here, we used an *in-vitro* ecosystem based on a number of these key members to test the hypotheses that beer ethanol content and bitterness (iso-α-acid level) affect the density and composition of the microbial community in the maturing beer, the beer chemistry, and the interactions between the microorganisms, the beer and the wood. To this end, a commercially available beer was used and its ethanol and iso-α-acid levels were adjusted from a ‘low’ level to a ‘medium’ and ‘high’ level. Microbial community dynamics were monitored at several time points via specific quantitative real-time PCR (qPCR) assays for each strain in the synthetic community, and beer chemistry was assessed at the same time points.

## 2 Materials and methods

### 2.1 Model system

To mimic the conditions of industrial-scale barrel-aging of beer, the model system displayed in Fig.1 was used. Briefly, the system was based on a 0.5-liter weck jar (diameter: 9.5 cm, height: 8.5 cm; Ikea, Zaventem, Belgium) in which the lid was replaced by a 3.5 cm thick wooden disk. The wood used represented new (unused) European oak (NIR category ‘Sweet’) from the cooperage ‘Garbellotto Spa’ (Pordenone, Italy) that would otherwise be used for barrel manufacturing. To mimic industrial maturation, the ratio between the wood contact surface and the beer volume was set to the same ratio as found in 225-liter barrels. Further, to ensure maximum contact between the beer and the wood, the jar was put upside down, while a rubber sealing ring between the wood and the jar was used to prevent leakage. Additionally, to ensure that oxygen ingress was not obstructed, jars were placed on a metal grid. Further, a water lock (diameter of the opening at the bottom: 9 mm; Brouwland, Beverlo, Belgium) was positioned through a 13-mm hole in the glass bottom of the jars using silicon plugs with a 9-mm hole for the water lock (Brouwland, Beverlo, Belgium) to allow pressure equalization and avoid pressure build-up inside the jar (Fig. 1). Before usage, the weck jars, rubber rings and water locks were thoroughly cleaned and disinfected with 70% ethanol. To saturize the wood pores and to extract the most pungent tannins, a pre-treatment was performed in which the jars were filled with a sterile solution of 0.1% citric acid (Vinoferm, Brouwland, Beverlo, Belgium) followed by an upside down incubation for five days at 20°C. The same pre-treatment is recommended when using new oak barrels in industrial barrel-aging (Bossaert *et al.*, 2021a; 2021b). Next, the jars were again disinfected with 70% ethanol, and filled with 500 ml filter-sterilized beer (see below).

**Figure 1:**
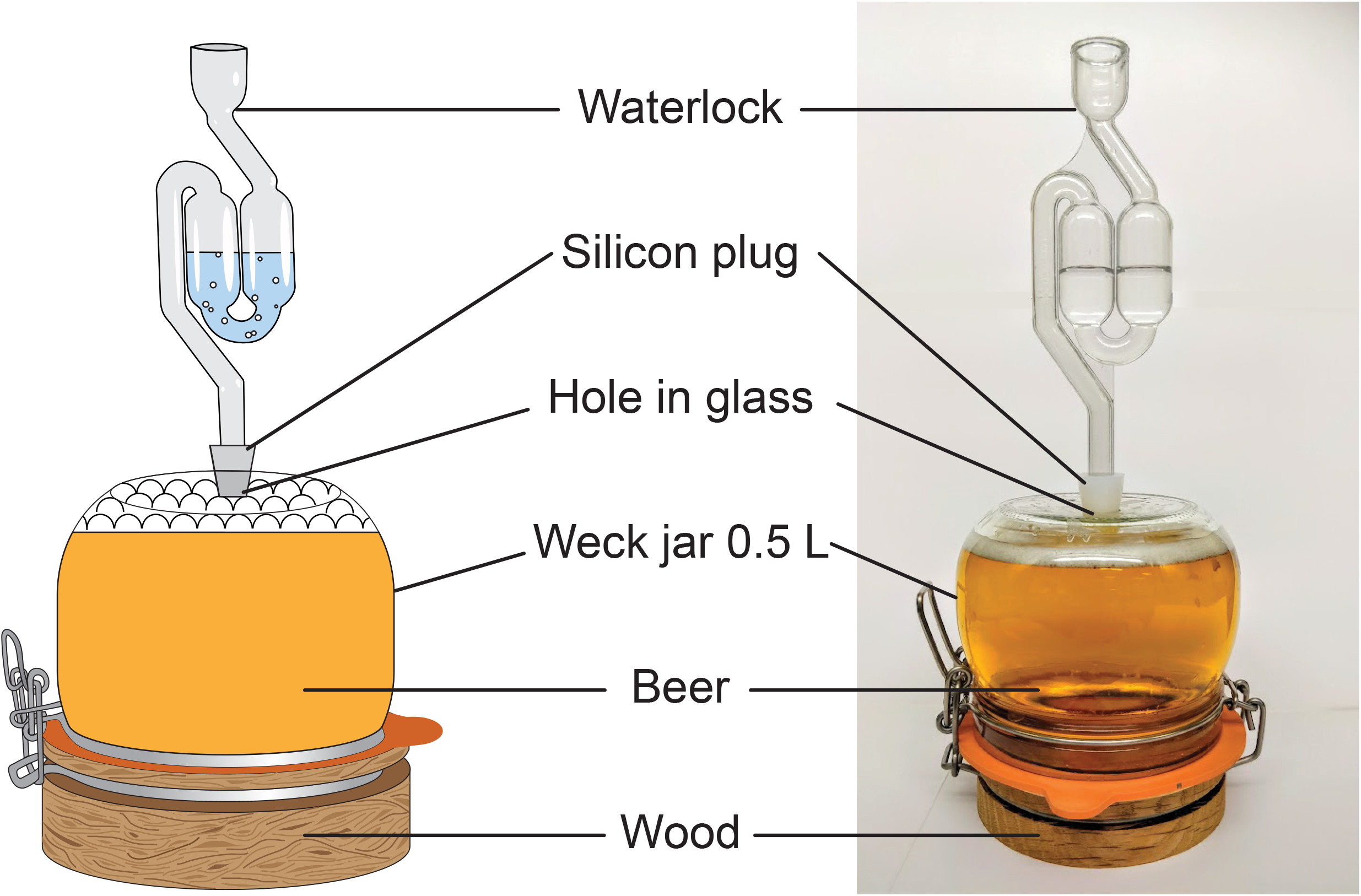
Illustration of the *in-vitro* system mimicking industrial barrel-aging on a lab scale. The design is based on a 0.5-liter weck jar in which the lid is replaced by a wooden disk made from wood that would otherwise be used for barrel manufacturing, i.e. new (unused) European oak. To ensure that the extraction of wood compounds and the ingress of oxygen is similar to industrial-scale barrel-aging, the ratio between the beer volume and the wood surface area is the same as in 225-liter barrels. A water lock is fitted through the hole in the bottom of the jar to avoid pressure build-up.

To simulate the microbial community in the aging process, a synthetic microbial community was assembled using eight key species that are consistently associated with barrel-aging of beer. The strains used were isolated previously from maturing beer (Bossaert *et al.*, 2021a) and represented members of the bacteria *Acetobacter malorum*, *Gluconobacter oxydans*, *Lactobacillus brevis* and *Pediococcus damnosus* (Table S1, Supplementary Information). Additionally, the community was composed of four strains belonging to the yeast species *Brettanomyces bruxellensis*, *Candida friedrichii*, *Pichia membranifaciens* and *Saccharomyces cerevisiae* (Table S1, Supplementary Information). In contrast to *G. oxydans* and *C. friedrichii* which, amongst several others, occur mainly at the beginning of the process, all other species have been found to dominate the microbial community over the course of maturation (Bossaert *et al.*, 2021a; 2021b; De Roos *et al.*, 2018). All strains were stored in 25% glycerol at −80°C.

To perform the experiments, strains were cultivated separately at 30°C on YPDF medium that contained 1% yeast extract (Oxoid, Thermo Fisher Scientific, Waltham, USA), 2% bactopeptone (BD, Franklin Lakes, USA), 1% D-glucose (Sigma-Aldrich, St. Louis, USA), 1% fructose (Acros Organics, Thermo Fisher Scientific, Waltham, USA) and 2% agar (Agar bacteriological no. 1, Oxoid, Thermo Fisher Scientific, Waltham, USA). All strains were incubated aerobically, with the exception of *L. brevis* and *P. damnosus*, which were cultivated in anaerobic conditions. Next, cells were collected, washed and resuspended in equal cell densities in sterile physiological water (0.85% NaCl; Merck, Darmstadt, Germany). Subsequently, all strains were combined and pitched to the beer (1 ml) at a final cell density of 10 colony forming units (CFU)/ml per strain in the beer. As a control, for every treatment (see below) jars were included in which the beer was not inoculated, but which also contained the wood. All jars were closed off by water locks filled with sterile water, and the weck jars were incubated statically in the dark at 20.0°C ± 1.1°C and a relative humidity of 52.7% ± 4.4% (Table S2, Supplementary Information). Under these conditions, preliminary experiments have shown that an incubation of 60 days led to similar microbial community dynamics as those observed during 38 weeks of industrial maturation (data not shown).

### 2.2 Treatments and sampling

The base beer used in this study (further called ‘reference beer’) was a Jupiler pilsner beer (Anheuser-Busch Inbev, Leuven, Belgium) with ‘low’ alcohol level (5.2 v/v%) and ‘low’ iso-α-acid level (13 ppm, corresponding to a bitterness of 17 international bitter units (IBU)) from one brewing batch. To assess the effect of ethanol, both a ‘medium’ and ‘high’ ethanol beer were prepared by adding 100% ethanol (Fisher Chemical, Thermo Fisher Scientific, Waltham, USA) to the reference beer to reach a final concentration of 8.0 v/v% and 11.0 v/v%, respectively. To test the effects of different iso-α-acid levels, 6% isomerized hop extract (Brewferm, Brouwland, Beverlo, Belgium) was added to the reference beer to obtain ‘medium’ (35 ppm, 50 IBU) and ‘high’ iso-α-acid levels (170 ppm, 135 IBU). Iso-α-acid concentrations were measured after filter sterilization by Ultra High Performance Liquid Chromatography (UPLC; protocol specifications provided in Table S3, Supplementary Information; the iso-α-acid composition of the beers is presented in Table S4, Supplementary Information). Samples were taken immediately after filling and pitching the jars (i.e. at day 0), and at day 10, 20, 30 and 60, by sacrificing three jars per treatment per time point (i.e. three biological replicates). The three negative controls (without microbes) per treatment were analyzed at the end of the experiment, i.e. at day 60. Sampling was performed by thoroughly shaking the beer, followed by removing the water lock and draining the entire homogenized beer volume through the hole in the bottom of the jar. The collected beer was then centrifuged at 3,500 × g for 15 minutes at 4°C, and obtained cell pellets and supernatants were preserved at −20°C for microbiological and chemical analyses, respectively.

### 2.3 Microbiological analyses

For every treatment and at each time point, inoculated strains were monitored by qPCR. DNA extraction was performed on 500 μl of the cell pellet (*n* = 2) according to the protocol described by Lievens *et al.* (2003). After combining both DNA replicates, qPCR amplification was performed (in duplicate) using species-specific primers and PrimeTime™ double-quenched 5’ 6-FAM/ZEN/3’ IBFQ probes (IDT, Leuven, Belgium) (Table S1, Supplementary Information). qPCR amplifications were performed according to the manufacturer’s protocol supplied with the PrimeTime® Gene Expression Master Mix (IDT, Leuven, Belgium) in a StepOnePlus Real-Time PCR system (Applied Biosystems, Thermo Fisher Scientific, Waltham, USA) using an annealing temperature of 62°C for all assays. In each qPCR run, a negative control in which template DNA was replaced by sterile DNA-free water was included. In order to estimate strain population densities, bacterial 16S ribosomal RNA (rRNA) gene and fungal Internal Transcribed Spacer (ITS) copy numbers were determined using standard curves based on a 10-fold amplicon dilution series for each strain. Further, to assess the total bacterial and fungal density, qPCR amplification was performed (in duplicate) using the bacterial 16S rRNA gene primers 515F and 806R (Caporaso *et al.*, 2011) and the fungal primers BITS and B58S3 (Bokulich and Mills, 2013) in combination with the iTaq™ Universal SYBR^®^ Green Supermix (Biorad Laboratories, Hercules, USA), as described previously (Bossaert *et al.*, 2021a). For these assays, gene copy numbers were determined using a standard 10-fold dilution of *A. malorum* and *S. cerevisiae* amplicons, respectively. The qPCR quantification cut-off was based on the obtained Ct-values of the qPCR negative controls and was set for all assays to a Ct-value of 34. For further data-analysis, log gene copy numbers were set to zero when the respective Ct-value was higher or equal to 34.

### 2.4 Chemical analyses

Chemical analyses were performed according to the methods described in Bossaert *et al.* (2021a). In total, eighteen aroma compounds related to wood maturation were measured via Headspace – Solid Phase Micro Extraction – Gas Chromatography – Mass Spectrometry (HS-SPME-GC-MS), and eighteen fermentation products (mainly higher alcohols and esters) were quantified via Headspace – Gas Chromatography – Flame Ionization Detector (HS-GC-FID). Five carbohydrates, three organic acids, pH, and total polyphenols were measured with a Gallery Plus Beermaster (Thermo Fisher Scientific, Waltham, USA), and the ethanol content was assessed using an Alcolyzer beer ME (Anton Paar GmbH, Graz, Austria). A detailed overview of the sample preparation protocols and appliance settings can be found in Table S3 (Supplementary Information). Further, for one weck jar that contained the inoculated reference beer, the concentration of dissolved oxygen in the beer was measured every ten minutes (from the beginning until the end of the experiment) using a FireSting-GO2 pocket oxygen meter with oxygen sensor spots applied to the jar (Pyro Science GmbH, Aachen, Germany).

### 2.5 Data visualization and statistical analyses

To test whether the microbial communities were significantly affected by the treatments and/or maturation time, permutational multivariate analysis of variance (perMANOVA) was performed on the qPCR data expressed as log gene copy numbers per μl DNA (Table S5, Supplementary Information) with the adonis function (vegan package) in R (v3.6.1), using 1,000 permutations (Oksanen *et al.*, 2019). Additionally, a non-metric multidimensional scaling (NMDS) plot was created with Bray-Curtis distances using the same data set to visualize differences in bacterial and fungal communities between treatments and maturation times, using the MDS function in R (vegan package, Oksanen *et al.*, 2019). Further, dynamics in microbial density are shown per strain as the average of the three biological replicates per treatment and time point. An additional perMANOVA with 1,000 permutations was performed with the adonis function of the vegan package in R (Oksanen *et al.*, 2019) to test for significant differences in beer chemistry across treatments and/or time points. In addition, principal component analysis (PCA) was applied to the scaled chemical data (stats package in R; R Core Team, 2019) to visualize differences in beer chemistry between all samples in a two-dimensional space. Moreover, for the most important chemical compounds, line plots were created using averages of the three biological replicates per treatment and time point. Finally, for each treatment and each chemical parameter the ratio of the compound in the inoculated beer and respective negative control beer at day 60 was calculated and visualized as a heatmap in MS Excel, while Welch t-tests were performed using the stats package in R, to test, for each treatment separately, whether the microbial community had a significant impact on the measured chemical parameters in comparison to the non-inoculated beers at day 60 (R Core Team, 2019). In like manner, to show the change in chemical parameters in the negative controls after 60 days of wood aging, ratios between the concentrations of chemical parameters in the negative controls at day 60 and the concentrations measured at day 0 were calculated, visualized in a heatmap, and subjected to Welch t-tests.

## 3 Results

### 3.1 Bacterial and fungal communities

Pitched bacteria and fungi were found in all jars inoculated with the synthetic microbial community, while they were not found in any of the negative controls, nor were other bacteria or fungi (as assessed by the total bacterial and fungal qPCRs and by plating). PerMANOVA revealed that treatment and maturation time significantly affected the bacterial and fungal community composition. Likewise, there was a significant interaction effect between both factors (Table 1). Additionally, the bacterial and fungal community composition in the three treatments with different ethanol levels was significantly different from each other, except for the fungal communities of the reference beer and the beer with a medium ethanol level. For the iso-α-acid treatments, microbial community composition differed between the reference beer and the beer with medium or high iso-α-acid levels (Table 1). Also, NMDS ordination of the qPCR data (stress bacteria = 0.067; stress fungi = 0.070) shows differences in the bacterial and fungal communities, especially across the different time points (represented by the first NMDS axis) (Fig. 2). *Lactobacillus brevis*, *P. damnosus*, and *C. friedrichii* were mainly associated with samples from day 0, while samples taken at a later time point contained more *A. malorum*, *G. oxydans*, *B. bruxellensis*, *P. membranifaciens*, and *S. cerevisiae*. Strongest effects on the bacteria can be observed for the ethanol treatments, for which the samples were separated the furthest from the samples of the reference beer, while the bacterial community composition in the iso-α-acid treatments remained more similar to the reference beer (Fig. 2A). In contrast, fungal communities in the different treatments converged to a similar community composition over the course of maturation, containing high cell densities of *S. cerevisiae*, *B. bruxellensis* and *P. membranifaciens* (Fig. 2B).

**Table 1:** Results of permutational multivariate analysis of variance (perMANOVA)^a^ comparing the bacterial and fungal community composition across the different ethanol (low level: 5.2 v/v%, medium level: 8 v/v%, high level: 11 v/v%) and iso-α-acid treatments (low level: 13 ppm and 17 IBU, medium level: 35 ppm and 50 IBU, high level: 170 ppm and 135 IBU), and/or different time points throughout a 60-day maturation^b^ of beer in a model system pitched with four bacterial^c^ and four fungal strains^d^.

| All treatments and time points combined |  |  |  |  |  |  |  |  |
| --- | --- | --- | --- | --- | --- | --- | --- | --- |
| Bacteria |  |  |  |  | Fungi |  |  |  |
| Factors | df | F | p |  | df | F | p |  |
| Treatment | 4 | 61.084 | < 0.001 | *** | 4 | 13.181 | < 0.001 | *** |
| Time | 4 | 67.390 | < 0.001 | *** | 4 | 77.058 | < 0.001 | *** |
| Treatment*time | 16 | 16.420 | < 0.001 | *** | 16 | 4.553 | < 0.001 | *** |
| Reference beer versus medium ethanol treatment |  |  |  |  |  |  |  |  |
| Bacteria |  |  |  |  | Fungi |  |  |  |
| Factors | df | F | p |  | df | F | p |  |
| Treatment | 1 | 62.815 | < 0.001 | *** | 1 | 1.7154 | 0.164 | n.s. |
| Time | 4 | 41.954 | < 0.001 | *** | 4 | 29.8046 | < 0.001 | *** |
| Treatment*time | 4 | 5.463 | 0.002 | ** | 4 | 2.4695 | 0.047 | * |
| Reference beer versus high ethanol treatment |  |  |  |  |  |  |  |  |
| Bacteria |  |  |  |  | Fungi |  |  |  |
| Factors | df | F | p |  | df | F | p |  |
| Treatment | 1 | 161.756 | < 0.001 | *** | 1 | 17.150 | < 0.001 | *** |
| Time | 4 | 39.652 | < 0.001 | *** | 4 | 34.854 | < 0.001 | *** |
| Treatment*time | 4 | 33.418 | < 0.001 | *** | 4 | 8.514 | < 0.001 | *** |
| Medium ethanol treatment versus high ethanol treatment |  |  |  |  |  |  |  |  |
| Bacteria |  |  |  |  | Fungi |  |  |  |
| Factors | df | F | p |  | df | F | p |  |
| Treatment | 1 | 58.111 | < 0.001 | *** | 1 | 12.8959 | < 0.001 | *** |
| Time | 4 | 30.805 | < 0.001 | *** | 4 | 29.6678 | < 0.001 | *** |
| Treatment*time | 4 | 18.275 | < 0.001 | *** | 4 | 4.0563 | 0.003 | ** |
| Reference beer versus medium iso-α-acid treatment | | | | | | | | |
| Bacteria |  |  |  |  | Fungi |  |  |  |
| Factors | df | F | p |  | df | F | p |  |
| Treatment | 1 | 12.472 | < 0.001 | *** | 1 | 5.161 | 0.018 | * |
| Time | 4 | 79.406 | < 0.001 | *** | 4 | 40.034 | < 0.001 | *** |
| Treatment*time | 4 | 3.806 | 0.008 | ** | 4 | 0.683 | 0.712 | n.s. |
| Reference beer versus high iso-α-acid treatment | | | | | | | | |
| Bacteria |  |  |  |  | Fungi |  |  |  |
| Factors | df | F | p |  | df | F | p |  |
| Treatment | 1 | 2.888 | 0.090 | n.s. | 1 | 12.011 | < 0.001 | *** |
| Time | 4 | 46.459 | < 0.001 | *** | 4 | 40.954 | < 0.001 | *** |
| Treatment*time | 4 | 8.462 | < 0.001 | *** | 4 | 2.147 | 0.055 | n.s. |
| Medium iso-α-acid treatment versus high iso-α-acid treatment | | | | | | | | |
| Bacteria |  |  |  |  | Fungi |  |  |  |
| Factors | df | F | p |  | df | F | p |  |
| Treatment | 1 | 0.871 | 0.379 | n.s. | 1 | 2.096 | 0.150 | n.s. |
| Time | 4 | 45.308 | < 0.001 | *** | 4 | 40.709 | < 0.001 | *** |
| Treatment*time | 4 | 2.417 | 0.077 | n.s. | 4 | 1.095 | 0.433 | n.s. |

<sup>a</sup> df: degrees of freedom, F: F statistic, p: p-value based on 1,000 permutations (values are statistically significant when p < 0.05), n.s.: not significant, \* p < 0.05, \*\* p < 0.01, \*\*\* p < 0.001.
<sup>b</sup> Beer samples were taken at the start (day 0) and after 10, 20, 30, and 60 days of wood maturation in the model system.
<sup>c</sup> Bacterial strains represented strains from *Acetobacter malorum*, *Gluconobacter oxydans*, *Lactobacillus brevis*, and *Pediococcus damnosus*.
<sup>d</sup> Fungal strains represented strains from *Brettanomyces bruxellensis*, *Candida friedrichii*, *Pichia membranifaciens*, and *Saccharomyces cerevisiae*.

**Figure 2:**
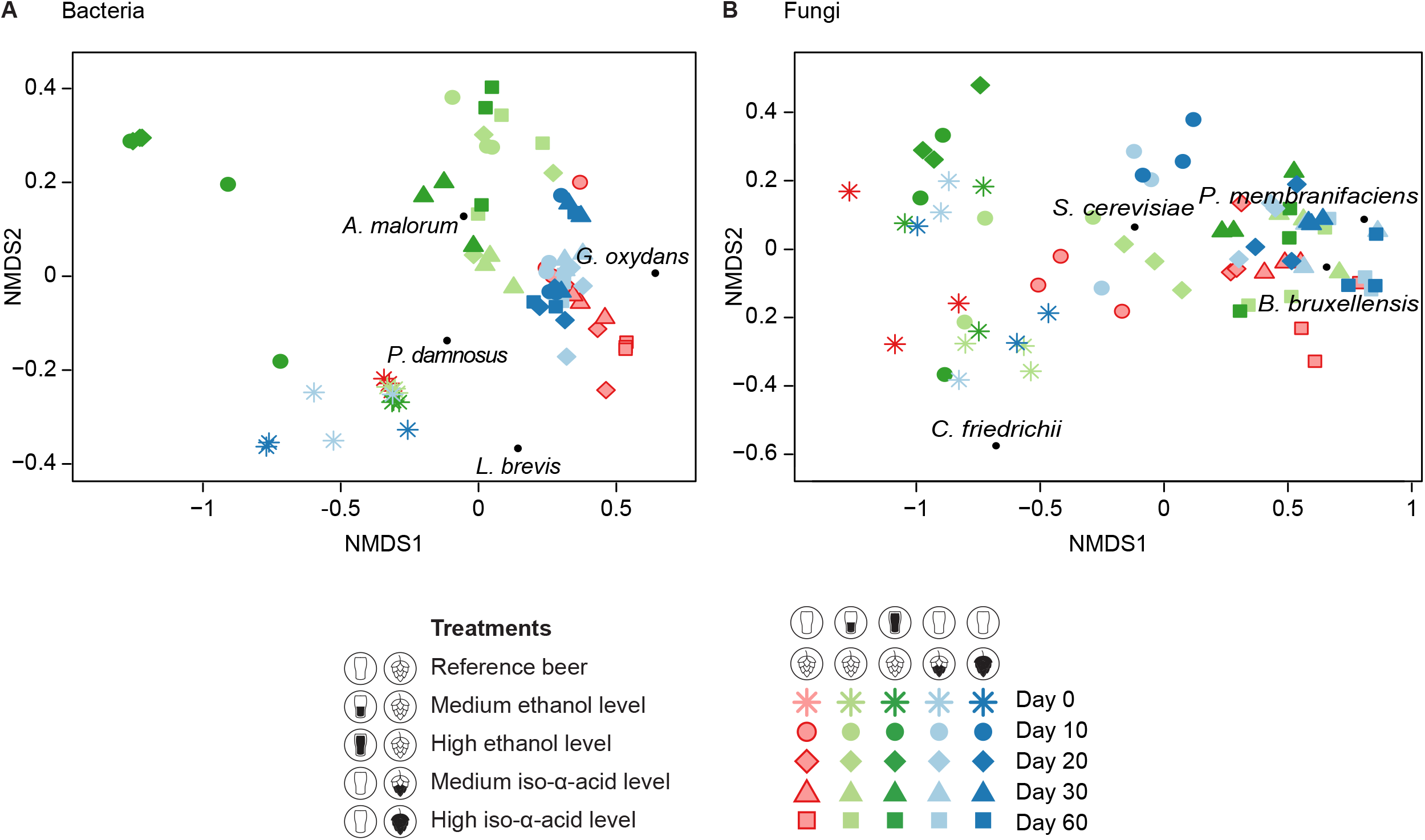
Non-metric multidimensional scaling (NMDS) ordination plots, based on Bray-Curtis distances of qPCR data (log gene copy numbers per μl DNA), visualizing differences in (A) bacterial (stress = 0.067) and (B) fungal (stress = 0.070) community composition of beer samples taken at different time points (indicated by different symbols) throughout a 60-day maturation period of beer in a model system in which a synthetic microbial community was pitched. The pitched community included four bacterial strains belonging to *Acetobacter malorum*, *Gluconobacter oxydans*, *Lactobacillus brevis* and *Pediococcus damnosus*, and four fungal strains belonging to *Brettanomyces bruxellensis*, *Candida friedrichii*, *Pichia membranifaciens* and *Saccharomyces cerevisiae*. In total, five treatments were investigated using three biological replicates (indicated by different colors): reference beer (low ethanol level: 5.5 v/v% and low iso-α-acid level: 13 ppm and 17 IBU), beer with medium ethanol level (8 v/v%), beer with high ethanol level (11 v/v%), beer with medium iso-α-acid level (35 ppm and 50 IBU), and beer with high iso-α-acid level (170 ppm and 135 IBU). The smaller the difference between two data points, the more similar the microbial communities. Microbial species are displayed in the NMDS plots as black dots.

Similar trends were observed when looking at the dynamics in cell densities of the total bacterial or total fungal populations and the cell densities of each strain separately (Fig. 3 and 4; Fig. S1, Supplementary Information). With the exception of the treatment with a high ethanol level, total bacterial density converged to a similar level for all treatments after 10 days of maturation (Fig. 3A-B). High ethanol treatment retarded bacterial growth, but total bacterial cell density converged to levels similar to the other treatments after 30 days of maturation (Fig. 3A). When zooming in on the individual strains, it becomes clear that not all strains contributed equally to the bacterial community structure. More specifically, *L. brevis* reached 7.30 ± 0.06 log 16S rRNA gene copy numbers per μl DNA in the reference beer after 60 days, while its growth was impeded by all ethanol and iso-α-acid treatments (reaching an average of 1.09 ± 0.28 log gene copy numbers after 60 days of maturation) (Fig. 3C-D). *Pediococcus damnosus* and *A. malorum* experienced a longer lag phase in the high ethanol treatment compared to the other treatments, but grew to similar cell densities for all treatments after 60 days of maturation, reaching 1.70 ± 0.11 and 7.09 ± 0.13 log 16S rRNA gene copy numbers per μl DNA, respectively (Fig. 3E-H). *Gluconobacter oxydans*, on the other hand, was not able to reach high cell densities in beers with a medium (2.58 ± 0.49 log 16S rRNA gene copy numbers) or high ethanol content (1.76 ± 0.09 log 16S rRNA gene copy numbers) in comparison to the reference beer (6.29 ± 0.07 log 16S rRNA gene copy numbers), but remained unaffected in the iso-α-acid treatments (medium level: 6.37 ± 0.13, high level: 5.77 ± 0.28 log 16S rRNA gene copy numbers) (Fig. 3I-J). In contrast, differences in cell densities between treatments were not as clear for the fungi (Fig. 4). Increasing the beer’s ethanol level mainly caused a longer lag phase of the fungi (Fig. 4A, 4C, 4E, 4G). On the contrary, addition of iso-α-acids did not affect fungal growth, except for *P. membranifaciens* (Fig. 4B, 4D, 4F, 4H, 4J). For the latter species, addition of iso-α-acids was found to stimulate fungal growth (up to 7.43 ± 0.56 log ITS copy numbers after 60 days of maturation in the high iso-α-acid treatment, opposed to 3.24 ± 1.94 log ITS copy numbers in the reference beer; Fig. 4H), and after 30 days of wood aging, ITS copy numbers of *P. membranifaciens* were higher in the medium iso-α-acid treatment than in the other treatments. Strikingly, *C. friedrichii* ITS copy numbers remained low and constant in all treatments, suggesting that this strain could not or only barely grow in the media tested (Fig. 4I-J).

**Figure 3:**
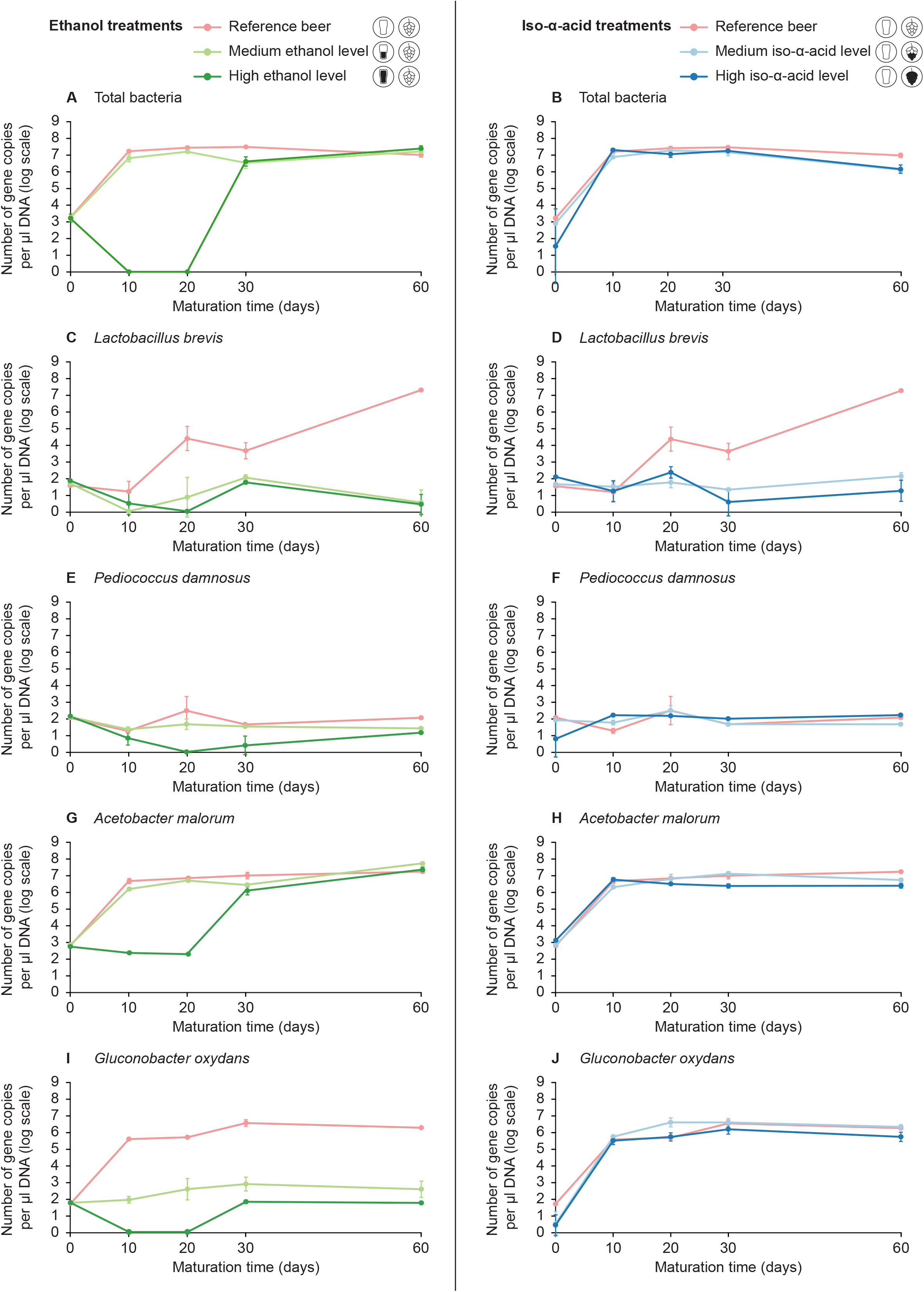
Temporal dynamics in 16S rRNA gene copy numbers for (A-B) the total bacterial community, (C-D) *Lactobacillus brevis*, (E-F) *Pediococcus damnosus*, (G-H) *Acetobacter malorum*, and (I-J) *Gluconobacter oxydans* throughout a 60-day maturation period of beer in a model system in which four bacteria (different panels) and four fungi (*Brettanomyces bruxellensis*, *Candida friedrichii*, *Pichia membranifaciens*, and *Saccharomyces cerevisiae*) were pitched. In total, five treatments were investigated (indicated by different colors): reference beer (low ethanol level: 5.5 v/v% and low iso-α-acid level: 13 ppm and 17 IBU), beer with medium ethanol level (8 v/v%), beer with high ethanol level (11 v/v%), beer with medium iso-α-acid level (35 ppm and 50 IBU), and beer with high iso-α-acid level (170 ppm and 135 IBU). Data are presented as the average of three biological replicates and the error bars represent the associated standard error of the mean.

**Figure 4:**
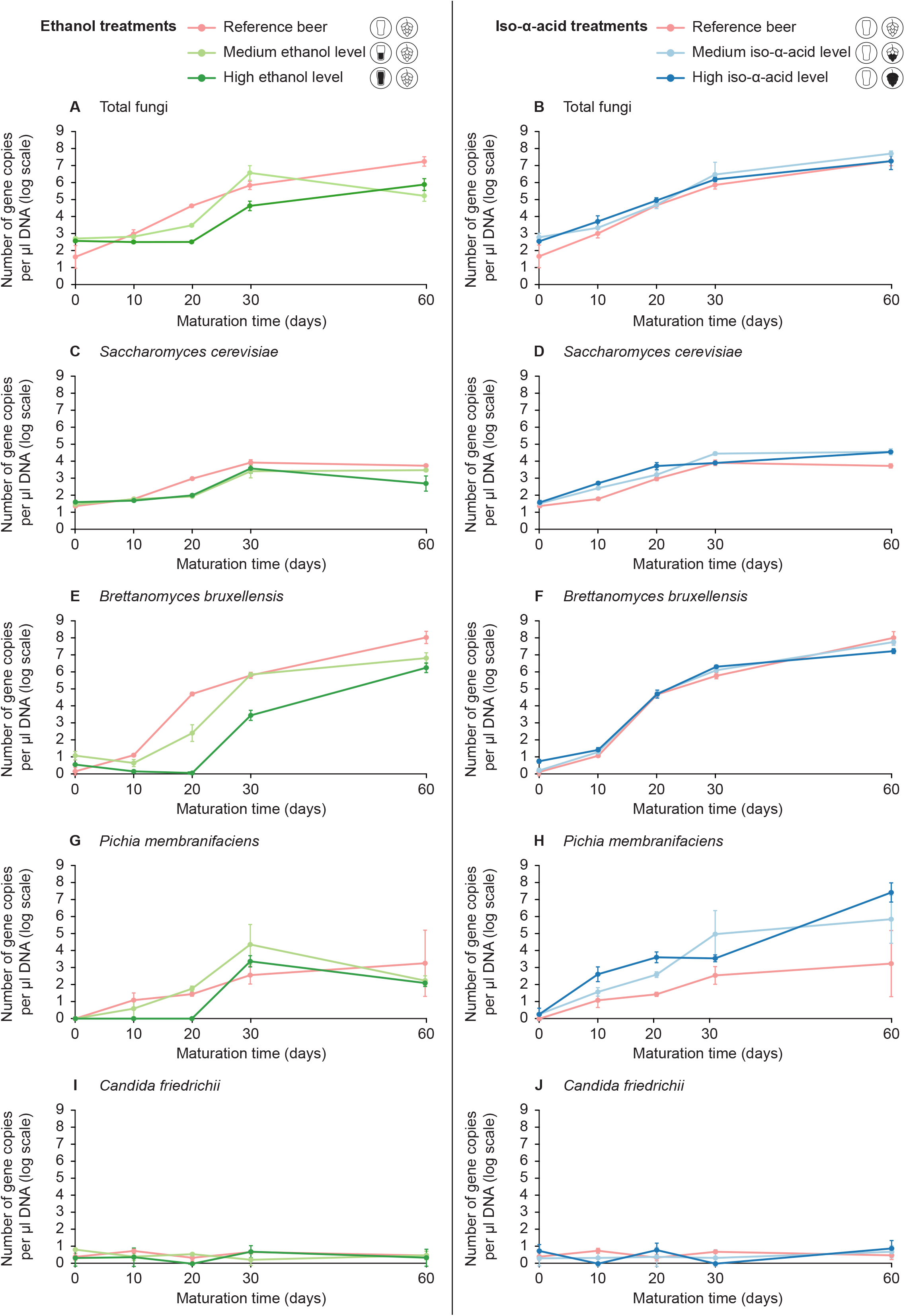
Temporal dynamics in ITS gene copy numbers for (A-B) the total fungal community, (C-D) *Saccharomyces cerevisiae*, (E-F) *Brettanomyces bruxellensis*, (G-H) *Pichia membranifaciens*, and (I-J) *Candida friedrichii* throughout a 60-day maturation period of beer in a model system in which four bacteria (*Acetobacter malorum*, *Gluconobacter oxydans*, *Lactobacillus brevis*, and *Pediococcus damnosus*) and four fungi (different panels) were pitched. In total, five treatments were investigated (indicated by different colors): reference beer (low ethanol level: 5.5 v/v% and low iso-α-acid level: 13 ppm and 17 IBU), beer with medium ethanol level (8 v/v%), beer with high ethanol level (11 v/v%), beer with medium iso-α-acid level (35 ppm and 50 IBU), and beer with high iso-α-acid level (170 ppm and 135 IBU). Data are presented as the average of three biological replicates and the error bars represent the associated standard error of the mean.

### 3.2 Beer chemistry

Measuring the dissolved oxygen concentration in the reference beer revealed that the oxygen concentration decreased within the first 59 hours from 1,806 ppb to 0 ppb, and remained 0 ppb until the end of the experiment (Table S6, Supplementary Information). PerMANOVA indicated that beer chemistry was significantly affected by the different treatments as well as by the maturation time. The interaction between both factors was also significant (Table 2). Moreover, beer samples with low, medium and high levels of either the ethanol or iso-α-acid treatments were significantly different from one another (Table 2). Both factors are also clearly separated in the PCA plot (Fig. 5). Samples taken at the different time points are distributed across the first diagonal (top left to bottom right corner), while samples of the different treatments are spread across the other diagonal (bottom left to top right corner) (Fig. 5). Furthermore, whereas samples from the ethanol treatments can be distinguished from other treatments quite clearly, the reference beer samples overlapped to some degree with the samples of the medium and high iso-α-acid treatments (Fig. 5), indicating that the chemical composition of the samples of the iso-α-acid treatments was more similar to the reference beer’s chemistry than to samples of the ethanol treatments. For all treatments, the pH decreased from 4.3 ± 0.0 to 3.9 ± 0.0, except for the high ethanol treatment where a pH of 4.0 ± 0.0 was reached after 60 days of maturation (Fig. 6A). For all treatments, a decrease in pH was accompanied by an increasing acetic acid concentration, reaching an average concentration of 1.69 ± 0.23 g/l (Fig. 6B). Furthermore, the L-lactic acid concentration was found to increase in the reference beer, from 28.5 ± 1.3 mg/l at the start of the experiment to 238.4 ± 37.1 mg/l at the end of the experiment (Fig. 6C). Further, the total sugar concentration (defined as the sum of D-glucose, D-fructose and sucrose) decreased to almost 0 mg/l, with a slower decline in samples of the ethanol treatments (Fig. 6D; Table S7, Supplementary Information). The concentration of 4-vinyl guaiacol decreased from 329.6 ± 26.8 to 61.2 ± 6.8 ppb on average for all treatments, except for the high ethanol treatment where a concentration of 161.5 ± 31.4 ppb was attained after 60 days of maturation (Fig. 6E). The concentrations of 4-ethyl guaiacol and 4-ethyl phenol increased over time, and encountered a quicker and larger incline in samples of the reference beer (from 0.5 ± 0.1 to 129.2 ± 9.7 and from 0.3 ± 0.0 to 77.4 ± 5.6, respectively) and the iso-α-acid treatments (on average from 1.2 ± 0.5 to 62.5 ± 7.5 and from 3.3 ± 1.0 to 73.0 ± 3.9, respectively) compared to the ethanol treatments (on average from 2.9 ± 1.2 to 34.3 ± 9.7 and from 2.9 ± 0.8 to 35.2 ± 6.7, respectively) (Fig. 6F-G). Moreover, remarkably, at day 30 substantially higher concentrations of 4-ethyl guaiacol and 4-ethyl phenol were recorded in the medium iso-α-acid treatment compared to the other treatments. Also for cis-3-methyl-4-octanolide the highest concentration (171.4 ± 23.9 ppb) was observed in the reference beer after 60 days of maturation, whereas the lowest concentration was measured in the high ethanol treatment (16.1 ± 7.9 ppb) (Fig. 6H). In contrast, the concentrations of eugenol and total polyphenols at day 60 could not be distinguished as clearly for all treatments (Fig. 6I-J), while other wood-related compounds like syringol, iso-eugenol, 4-vinyl guaiacol, 4-methyl guaiacol, guaiacol, methyl vanillate and vanillin were found in higher concentrations in at least one of the ethanol treatments (Table S8, Supplementary Information). In fact, the concentration of vanillin reached a maximum value in all treatments before the end of the experiment. Highest maximum values were obtained for the ethanol treatments, reaching a maximum level of 336.4 ± 90.3 ppb after 10 days of maturation in the medium ethanol treatment and a maximum level of 528.1± 51.4 ppb after 30 days in the high ethanol treatment (Fig. 6K). A similar trend is observed for furfural and 5-methyl furfural, i.e. while concentrations at the beginning and at the end of the experiment were similarly low (57.4 ± 5.3 and 8.2 ± 1.7 ppb, respectively), highest furfural (2747.1 ± 671.0 ppb) and 5-methyl furfural (606.8 ± 247.3 ppb) concentrations were obtained in the high ethanol treatment after 20 and 30 days of maturation, respectively (Fig. 6L-M). Finally, the concentration of certain esters including ethyl acetate was found to increase over time, while other esters like isoamyl acetate experienced a decrease (Fig. 6N-O) (Table S7, Supplementary Information).

**Table 2:** Results of permutational multivariate analysis of variance (perMANOVA)^a^ comparing beer chemistry across the different ethanol (low level: 5.2 v/v%, medium level: 8 v/v%, high level: 11 v/v%) and iso-α-acid treatments (low level: 13 ppm and 17 IBU, medium level: 35 ppm and 50 IBU, high level: 170 ppm and 135 IBU), and/or different time points throughout a 60-day maturation^b^ of beer in a model system pitched with four bacterial^c^ and four fungal strains^d^.

| All treatments and time points combined |  |  |  |  |
| --- | --- | --- | --- | --- |
| Factors | df | *F* | *p* |  |
| Treatment | 4 | 16.0358 | < 0.001 | *** |
| Time | 4 | 18.7659 | < 0.001 | *** |
| Treatment*time | 16 | 4.2046 | < 0.001 | *** |
| Reference beer versus medium ethanol treatment |  |  |  |  |
| Factors | df | *F* | *p* |  |
| Treatment | 1 | 8.8857 | < 0.001 | *** |
| Time | 4 | 18.8185 | < 0.001 | *** |
| Treatment*time | 4 | 3.5766 | < 0.001 | *** |
| Reference beer versus high ethanol treatment |  |  |  |  |
| Factors | df | *F* | *p* |  |
| Treatment | 1 | 26.2434 | < 0.001 | *** |
| Time | 4 | 9.0025 | < 0.001 | *** |
| Treatment*time | 4 | 6.0169 | < 0.001 | *** |
| Medium ethanol treatment versus high ethanol treatment |  |  |  |  |
| Factors | df | *F* | *p* |  |
| Treatment | 1 | 18.7839 | < 0.001 | *** |
| Time | 4 | 8.9067 | < 0.001 | *** |
| Treatment*time | 4 | 5.3771 | < 0.001 | *** |
| Reference beer versus medium iso-α-acid treatment | | | | |
| Factors | df | *F* | *p* |  |
| Treatment | 1 | 4.4242 | 0.006 | ** |
| Time | 4 | 14.4355 | < 0.001 | *** |
| Treatment*time | 4 | 2.5080 | 0.003 | ** |
| Reference beer versus high iso-α-acid treatment | | | | |
| Factors | df | *F* | *p* |  |
| Treatment | 1 | 6.3957 | < 0.001 | *** |
| Time | 4 | 12.9780 | < 0.001 | *** |
| Treatment*time | 4 | 2.9596 | 0.002 | ** |
| Medium iso-α-acid treatment versus high iso-α-acid treatment | | | | |
| Factors | df | *F* | *p* |  |
| Treatment | 1 | 3.3800 | 0.020 | * |
| Time | 4 | 8.5526 | < 0.001 | *** |
| Treatment*time | 4 | 1.9624 | 0.027 | * |

<sup>a</sup> df: degrees of freedom, *F*: F statistic, *p*: p-value based on 1,000 permutations (values are statistically significant when $p < 0.05$ ), n.s.: not significant, \* $p < 0.05$ ; \*\* $p < 0.01$ ; \*\*\* $p < 0.001$ .
<sup>b</sup> Beer samples were taken at the start (day 0) and after 10, 20, 30, and 60 days of wood maturation in the model system.
<sup>c</sup> Bacterial strains represented strains from *Acetobacter malorum*, *Gluconobacter oxydans*, *Lactobacillus brevis*, and *Pediococcus damnosus*.
<sup>d</sup> Fungal strains represented strains from *Brettanomyces bruxellensis*, *Candida friedrichii*, *Pichia membranifaciens*, and *Saccharomyces cerevisiae*.

**Figure 5:**
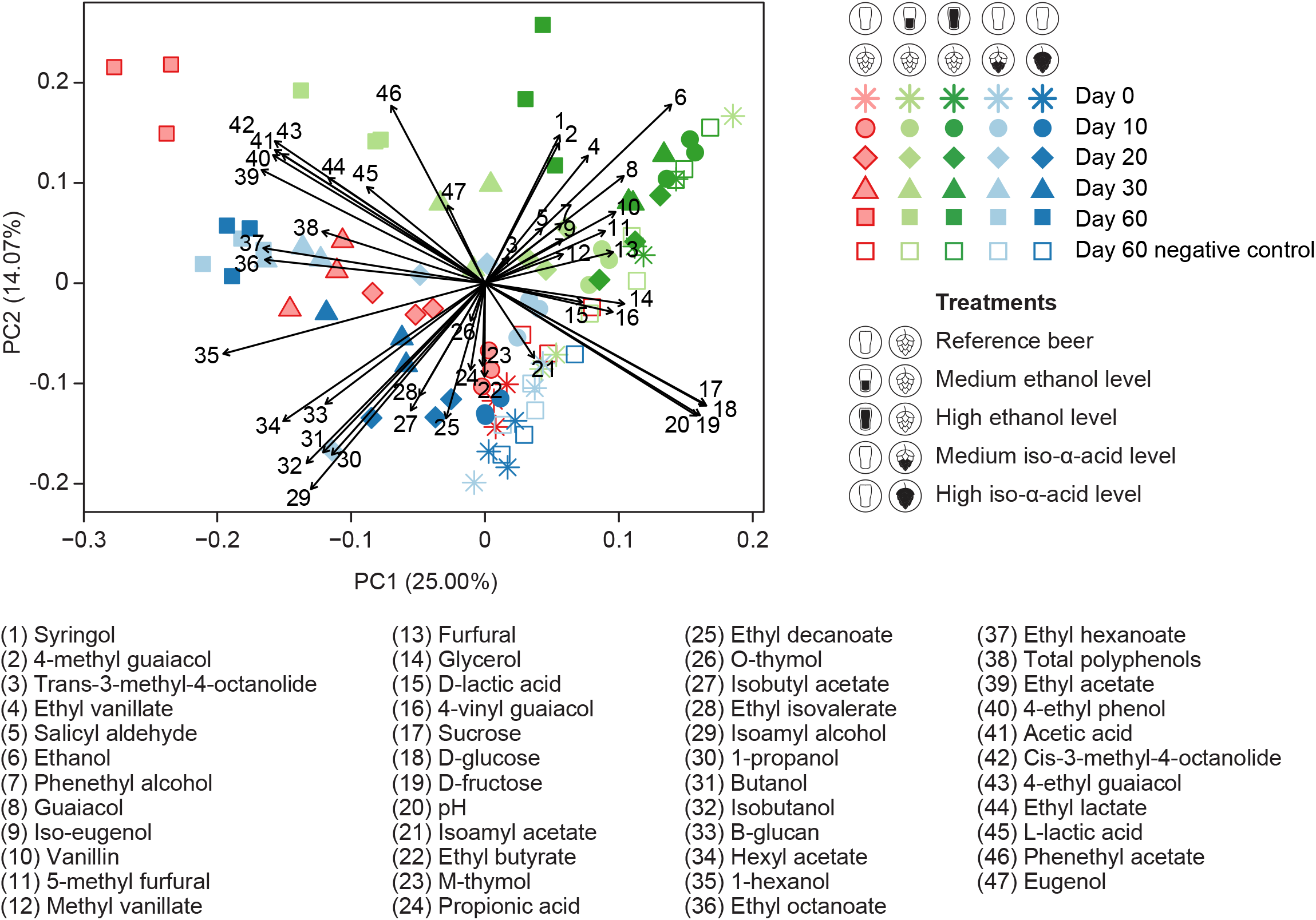
Principal component analysis (PCA) visualizing the differences in chemical composition of beer samples taken at different time points (indicated by different symbols) throughout a 60-day maturation period of beer in a model system in which a synthetic microbial community was pitched. The pitched community included four bacterial strains belonging to *Acetobacter malorum*, *Gluconobacter oxydans*, *Lactobacillus brevis* and *Pediococcus damnosus*, and four fungal strains belonging to *Brettanomyces bruxellensis*, *Candida friedrichii*, *Pichia membranifaciens* and *Saccharomyces cerevisiae*. In total, five treatments were investigated using three biological replicates (indicated by different colors): reference beer (low ethanol level: 5.5 v/v% and low iso-α-acid level: 13 ppm and 17 IBU), beer with medium ethanol level (8 v/v%), beer with high ethanol level (11 v/v%), beer with medium iso-α-acid level (35 ppm and 50 IBU), and beer with high iso-α-acid level (170 ppm and 135 IBU). Negative controls with wood, but without microbial community, were analyzed after 60 days of maturation and are plotted as open circles for each treatment. The chemical variables are presented as vectors. The smaller the difference between two data points, the more similar the chemical composition.

**Figure 6:**
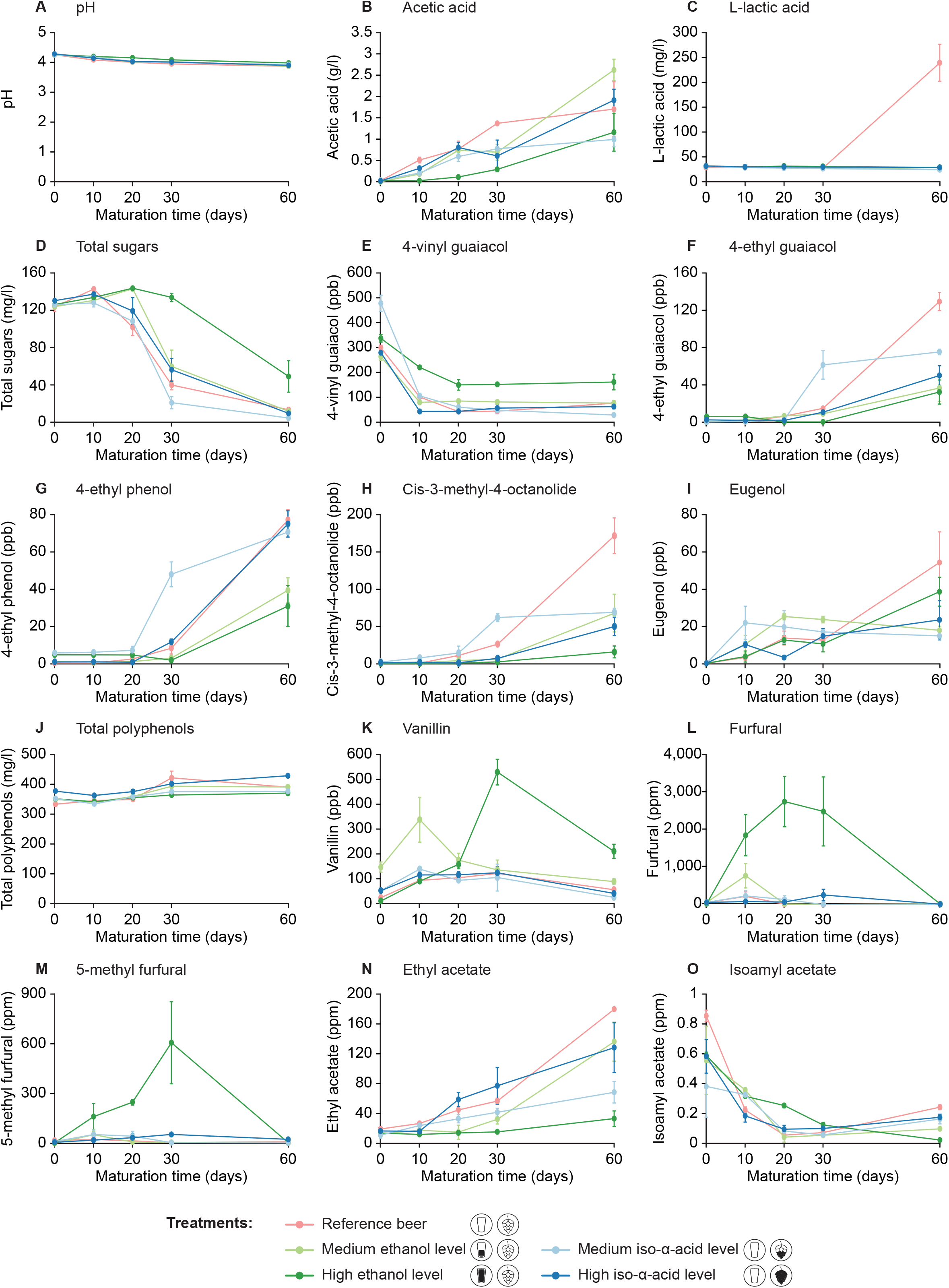
Temporal changes in beer chemistry throughout a 60-day maturation period of beer in a model system in which a synthetic microbial community was pitched. The pitched community included four bacterial strains belonging to *Acetobacter malorum*, *Gluconobacter oxydans*, *Lactobacillus brevis* and *Pediococcus damnosus*, and four fungal strains belonging to *Brettanomyces bruxellensis*, *Candida friedrichii*, *Pichia membranifaciens* and *Saccharomyces cerevisiae*. In total, five treatments were investigated (indicated by different colors): reference beer (low ethanol level: 5.5 v/v% and low iso-α-acid level: 13 ppm and 17 IBU), beer with medium ethanol level (8 v/v%), beer with high ethanol level (11 v/v%), beer with medium iso-α-acid level (35 ppm and 50 IBU), and beer with high iso-α-acid level (170 ppm and 135 IBU). Data are presented as the average of three biological controls and the error bars represent the associated standard error of the mean. Displayed parameters: (A) pH, (B) acetic acid, (C) L-lactic acid, (D) total sugars (defined as the sum of D-glucose, D-fructose and sucrose), (E) 4-vinyl guaiacol, (F) 4-ethyl guaiacol, (G) 4-ethyl phenol, (H) cis-3-methyl-4-octanolide (cis-oak lactone), (I) eugenol, (J) total polyphenols, (K) vanillin, (L) ethyl acetate, (M) isoamyl acetate, (N) furfural, and (O) 5-methyl furfural. For a detailed overview of the different chemical parameters measured in this study, the reader is referred to Tables S7 and S8 (Supplementary Information).

When zooming in on the chemical profiles obtained in the negative controls after 60 days of wood maturation, the concentrations of many compounds were found to be significantly different from the concentrations measured at day 0 (Fig. S2, Supplementary Information). Concentrations of D-glucose, D-fructose and sucrose were significantly higher in the negative controls of most treatments after 60 days of wood aging, whereas the concentrations of 4-vinyl guaiacol, and certain esters like ethyl butyrate, ethyl hexanoate, ethyl octanoate and isoamyl acetate were significantly lower than the concentrations measured at day 0 (Fig. S2, Supplementary Information). Further, although not (consistently) significant, the concentrations of certain wood compounds including cis- and trans-3-methyl-4-octanolide, vanillin, furfural and 5-methyl furfural were higher in the negative controls at day 60 than at the start of the maturation (Fig. S2, Supplementary Information). Likewise, in comparison with the negative controls at day 60, some compounds are found in substantially lower concentrations at day 60 in jars where microorganisms were pitched (Fig. 7), indicating that these compounds were utilized or converted by the microbes. More specifically, the concentrations of the following compounds were at least ten times lower in the inoculated samples than in the negative controls: D-glucose, D-fructose, sucrose, furfural, and 5-methyl furfural (Fig. 7). Additionally, for four out of five treatments, the concentrations of vanillin at day 60 were more than 50% lower in samples with a microbial community (Fig. 7). Further, concentrations of eugenol and iso-eugenol were also lower in inoculated samples, although the differences in concentration were less outspoken. In contrast, concentrations of acetic acid, 4-ethyl guaiacol, 4-ethyl phenol, cis-3-methyl-4-octanolide, ethyl acetate, ethyl lactate, ethyl hexanoate, and phenethyl acetate were more than ten times higher in inoculated samples of some of the treatments (Fig. 7). Overall, in the reference beer, 23 of the 42 chemical compounds measured could be significantly linked to the microbial community, while in the medium ethanol and high iso-α-acid treatments 19 compounds were significantly linked to the presence of microbes, followed by 16 compounds in the medium iso-α-acid treatment, and 7 compounds in the high ethanol (Fig. 7).

**Figure 7:**
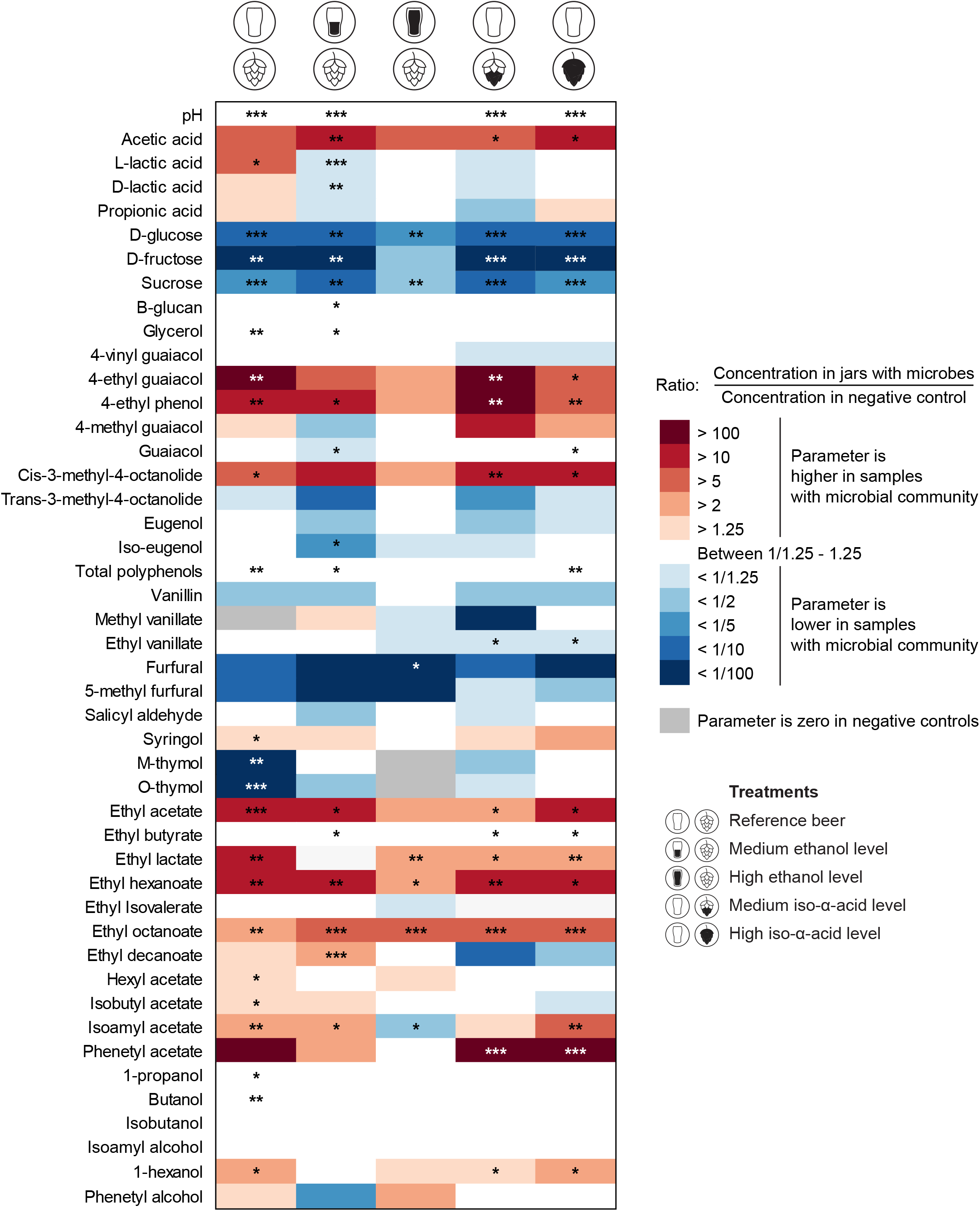
Heatmap visualizing the ratio between the concentration of the different chemical compounds in the jars spiked with a synthetic microbial community and the negative controls without microbes, indicating whether compound concentrations were higher (red) or lower (blue) in the inoculated jars compared to the respective negative controls. The pitched community included four bacterial strains belonging to *Acetobacter malorum*, *Gluconobacter oxydans*, *Lactobacillus brevis* and *Pediococcus damnosus*, and four fungal strains belonging to *Brettanomyces bruxellensis*, *Candida friedrichii*, *Pichia membranifaciens* and *Saccharomyces cerevisiae*. Results are presented for the five different treatments (*n* = 3), including: reference beer (low ethanol level: 5.5 v/v% and low iso-α-acid level: 13 ppm and 17 IBU), beer with medium ethanol level (8 v/v%), beer with high ethanol level (11 v/v%), beer with medium iso-α-acid level (35 ppm and 50 IBU), and beer with high iso-α-acid level (170 ppm and 135 IBU). Welch t-tests were performed to test whether the effect of the microbial community on the chemical parameters measured was significant. *P*-values obtained in the t-tests are presented in the cells: *\* p* < 0.05, ** *p* < 0.01, *** *p* < 0.001.

## 4 Discussion

Regardless of treatments, total bacterial and total fungal cell densities increased substantially throughout maturation, up to densities corresponding to 7.4 and 7.7 log gene copy numbers, respectively. These results were confirmed by plating subsamples on a number of cultivation media, reinforcing the robustness of our results (Table S9; Fig. S3, Supplementary Information). Nevertheless, densities of individual strains were largely dependent on the treatments applied, resulting in different microbial community dynamics over the course of maturation among treatments, especially for bacteria. For example, growth of *L. brevis* and *G. oxydans* was inhibited up until the end of the experiment when ethanol concentrations were elevated, while the other bacterial strains could still reach cell densities similar to those obtained in the reference beer (5.2 v/v% alcohol). Nevertheless, although similar densities were obtained, growth of *P. damnosus* and *A. malorum* was substantially delayed in the high ethanol beer (11 v/v%), suggesting a strong impact of ethanol on the lag phase period. By contrast, fungal community composition at day 60 was more similar for all treatments, even though ethanol affected fungal growth and increased the lag phase for all fungal strains. The lag phase of *B. bruxellensis* and *P. membranifaciens* in the medium ethanol treatment was longer than in the reference beer, yet shorter than in the high ethanol treatment, while for *S. cerevisiae* the lag phases in both the medium and high ethanol treatment were comparable, yet longer than in the reference beer. These findings are in agreement with Chandra *et al.* (2014), showing an increase in the lag phase of pure-culture *Brettanomyces* spp. in media with higher ethanol concentrations. Altogether, these results suggest that *P. damnosus* and *A. malorum* might have the highest ethanol tolerance of all strains included in the synthetic community, followed by *B. bruxellensis, P. membranifaciens*, and *S. cerevisiae*, whereas *L. brevis* and *G. oxydans* are seemingly the most sensitive to ethanol. While it is well known that ethanol can affect microbial strains directly as it is a common stress factor for microbial growth (Haakensen *et al.*, 2009; Kato *et al.*, 2011), high ethanol concentrations can also affect microorganisms indirectly by prolonging the lag phase and providing an opportunity to other microbes to dominate the community (Barrajón *et al.*, 2010; García-Ríos *et al.*, 2014). In case the growth of certain strains depends on the presence of other strains (e.g. when they rely on other strains to supply nutrients), a prolonged lag phase of a given strain may also induce a longer lag phase of other strains that depend on the first strain (Morris *et al.*, 2013; Seth and Taga, 2014). As a result, the competition degree or fitness level of a strain may not only be affected by abiotic factors, but also by biotic factors like the presence of other microorganisms or killer toxins, which determine the capacity of one strain to out-compete another (Ciani *et al.*, 2016). More research focusing on the mechanisms of microbial community assembly could provide more insight into this matter.

Likewise, the iso-α-acid treatments affected microbial growth, in particular that of *L. brevis* and *P. membranifaciens*. *Lactobacillus brevis* was not able to grow to cell densities similar to those obtained in the reference beer with a low bitterness (13 ppm iso-α-acids), and its growth was equally restricted by both iso-α-acid concentrations tested (35 and 170 ppm). Indeed, it is generally known that hop iso-α-acids are able to penetrate the cell membrane of gram-positive bacteria like lactic acid bacteria, change the intracellular pH and disturb membrane-bound processes, leading to cell death (Sakamoto and Konings, 2003; Zhao *et al.*, 2017). Nevertheless, growth of the other lactic acid bacterium included, *P. damnosus*, was not affected by the tested iso-α-acid concentrations, most probably because the tested strain had better adapted to the beer environment, rendering it more resistant to iso-α-acids (Sakamoto and Konings, 2003). In contrast, *P. membranifaciens* grew to higher cell densities in the beers with elevated iso-α-acids concentrations in comparison to the reference beer. While it has been described that *Pichia* spp. can be involved in the biotransformation of hop terpenes (incl. geraniol) (Ponzoni *et al.*, 2008), it remains unclear whether *P. membranifaciens* can use hop-derived compounds from the iso-α-extracts as a substrate for growth.

In addition to the microbial community composition, beer chemistry was significantly affected by all treatments and by maturation time. Specifically, several of the wood-related compounds measured in this study (e.g. syringol, iso-eugenol, vanillin, methyl vanillate, guaiacol and 4-vinyl guaiacol) were found in higher concentrations in the beers with higher ethanol levels. These findings are in line with Sterckx *et al.* (2012b), demonstrating that a higher ethanol concentration induces a higher extraction rate of wood compounds as they generally dissolve better in ethanol than in water. Besides the chemical extraction of compounds from the wood, microbial activity may also strongly affect beer chemistry. More specifically, it appears that the microbes pitched in the beer consumed sugars at a rate that is proportional to their growth rate, and produced organic acids (causing a decrease in pH) and a wide variety of other metabolites over the course of maturation. In fact, L-lactic acid only reached high concentrations in the reference beer and was absent in the respective negative control, indicating that it was the result of microbial activity. Furthermore, as *L. brevis* was the only microorganism that was inhibited by the different ethanol and iso-α-acid treatments, the majority of lactic acid was most probably produced by *L. brevis*. Further, the decrease in 4-vinyl guaiacol and increase in 4-ethyl guaiacol suggests that a conversion reaction might have taken place, possibly due to the activity of microbial hydroxycinnamic acid decarboxylases (Dzialo *et al.*, 2017; Saez *et al.*, 2011). Indeed, the concentration of 4-ethyl guaiacol was much higher in the jars with microbes than in the negative controls, and is thus most probably linked to microbial activity. The latter observation is also confirmed by the associated increase in 4-ethyl phenol in the jars spiked with microbes, which is known to be produced from 4-vinyl phenol (Dzialo *et al.*, 2017). Moreover, as the concentration of 4-ethyl guaiacol in the reference beer at day 60 is considerably higher than in the other treatments, and as *L. brevis* was the only organism that could not thrive well under the applied conditions except in the reference beer, this may indicate that *L. brevis* has played a major role in the production of 4-ethyl guaiacol. This hypothesis is in agreement with previous findings showing that *L. brevis* can produce ethyl phenolic compounds from their vinyl phenolic precursors (Couto *et al.*, 2006). Nevertheless, it was also found that production of 4-ethyl guaiacol and 4-ethyl phenol depends largely on the environmental conditions (pH, precursors available, etc.) (Silva *et al.*, 2011). Furthermore, it was found that an increase in *Lactobacillus* biomass resulted in a higher production of ethyl phenolic compounds (Chandra *et al.*, 2014; Silva *et al.*, 2011), while Van Beek and Priest (2000) suggested that the production of ethyl phenolics by *Lactobacillus* is enhanced in the presence of *S. cerevisiae*. Strikingly, the concentrations of 4-ethyl guaiacol and 4-ethyl phenol at day 30 in the medium iso-α-acid treatment were higher than in the other treatments, an observation that aligns well with the growth profiles of some of the microbes included in the system, particularly *P. membranifaciens*, which is known to produce ethyl phenolic compounds (Saez *et al.*, 2011). Hence, this might be an indication that the *P. membranifaciens* strain used in this study is also involved in the generation of 4-ethyl guaiacol and 4-ethyl phenol. Nevertheless, also strains from other species used in this experiment may possess similar enzyme activities, including *B. bruxellensis* and some *Pediococcus* strains (Couto *et al.*, 2006; Steensels *et al.*, 2015).

Besides the consumption of carbohydrates from the beer and the production of common microbial metabolites, our results suggest that there is a more intricate interaction taking place between the microorganisms, the beer and the wood. Although low concentrations of sugars were measured in the beers at the beginning of the experiment, and the negative controls indicated that small amounts of sugars were directly extracted from the wood, enriching the beers with additional sugars, these concentrations would most probably not be sufficient to support microbial growth to the cell densities that were found at the end of the maturation (Gao *et al.*, 2021; Greenman *et al.*, 1981). This suggest that other carbon sources may come into play or that the microorganisms involved can generate accessible carbon sources from the wood. Indeed, besides the residual sugars in the beer, degradation products of wood cellulose and hemicellulose can be used as carbon sources for microbial growth (Gollihue *et al.*, 2018). Microorganisms like *B. bruxellensis* and certain strains of *P. membranifaciens* and *Candida* spp. exhibit β-glucosidase activity and can generate fermentable carbon sources from cellobiose, a degradation product of cellulose (Crauwels *et al.*, 2015; Genovés *et al.*, 2003; López *et al.*, 2015). Furthermore, *P. membranifaciens* can produce 1,4-β-xylosidase that plays a role in the hydrolysis of xylan, i.e. one of the major components of hemicelluloses found in the cell walls of monocots and hard woods (López *et al.*, 2015; Romero *et al.*, 2012). Additionally, specific wood compounds can have a beneficial effect on the growth of certain microbes, as demonstrated by de Revel *et al.* (2005). Although the mechanisms are still unclear, the authors showed that growth of the lactic acid bacterium *Œnococcus œni* was enhanced in the presence of vanillin. Fungal species, on the other hand, can be inhibited by high concentrations of vanillin (Fitzgerald *et al.*, 2003), and possibly for that reason, often possess the capability to convert vanillin into other phenolic derivatives like vanillyl alcohol (Fitzgerald *et al.*, 2003). In this study, vanillin was found to be extracted from the wood, as shown by the relatively high vanillin concentrations in the negative controls, while vanillin concentrations in the jars with the synthetic microbial community were much lower, suggesting a microbial conversion of this compound. Both yeasts and bacteria have been found to convert vanillin into other monophenolic compounds, including some of the species used in our experiments (Delgenès *et al.*, 1996 ; De Wulf *et al.*, 1987; Edlin *et al.*, 1995). A similar trend was observed for furfural and 5-methyl furfural, i.e. an extraction from the wood as shown by the negative controls, but a lower concentration in samples that contained a microbial community. This trend may be explained by the activity of *P. membranifaciens*, which shows a growth pattern in line with the dynamics in the furfural concentration. Moreover, previous research has shown that xylose reductase produced by *Pichia* spp. can reduce furfural and hydroxyl-methyl furfural concentrations (Almeida *et al.*, 2008). Additionally, lactic acid bacteria and *G. oxydans* have been associated with furfural degradation as well (Bastard *et al.*, 2016; Zhou *et al.*, 2017), but the growth profiles of these bacteria did not align with the dynamics in furfural concentration. Further, the concentration of the common wood-derived compound cis-3-methyl-4-octanolide, also called ‘cis-oak lactone’, remained rather low in the samples without pitched microorganisms, while much higher values were measured in the beer samples with microorganisms, especially in the reference beer. This indicates that microbial activity is most likely responsible for the increase in concentration of this compound. As *L. brevis* is the only species that is much more abundant in the reference beer than in all other treatments, *L. brevis* is likely involved in this reaction. Likewise, Bastard *et al.* (2016) have reported an increase in the extraction of oak lactones in the presence of lactic acid bacteria.

Altogether, our results show that an intricate interaction takes place between the wood, the microbes and the beer during wood-aging of beer. In fact, this is probably one of the main reasons why wood-aging of conventionally fermented beer largely remains a trial-and-error process and why there is often inconsistency and unpredictability in the product quality of wood-aged beverages. More research focusing on the interactions between wood, microbes and the maturing beer could help unravel the reactions that take place and could lead to the development of new strategies to improve process control and predictability. The use of model systems like the one used in this study, is an indispensable and cost-effective tool to study these interactions in-depth, one parameter at the time, and under controlled conditions (Wolfe and Dutton, 2015; Wolfe 2018). Furthermore, it allows to study more general ecological questions in-depth, including the study of microbial interactions and mechanisms of microbial community assembly. Further, the system allows to screen for new unconventional yeasts and bacteria that can be used to add additional flavors to wood-aged beers. Ultimately, the system may yield plenty of opportunities for beer diversification opening the door to the production of new, innovative beers with a variety of complex, wood-derived flavor profiles.

## Supporting information

Fig S1 Line plots per treatment

Fig S2 Chemistry heatmaps negative controls

Fig S3 Plate counts

## 7 Ethical guidelines

Ethics approval was not required for this research.

## 8 Supplementary Material

**Table S1**: Microbial strains used in this study, including primer and probe sequences used for qPCR analysis

**Table S2**: Data log of temperature, humidity and dew point over the course of the experiment

**Table S3**: Specifications of the chemical analysis protocols

**Table S4**: Composition of the iso-α-acid fraction in the different beer media tested

**Table S5**: qPCR data expressed as gene copy numbers per μl DNA (in log units)

**Table S6**: Data log of dissolved oxygen concentration in the reference beer inoculated with the synthetic microbial community

**Table S7**: pH and concentration of carbohydrates, ethanol and fermentation products

**Table S8**: Concentrations of wood and hop compounds

**Table S9**: Composition of cultivation media and incubation conditions

**Figure S1**: Temporal dynamics in bacterial (left panels) and fungal (right panels) 16S rRNA gene and ITS copy numbers, respectively, throughout the maturation process of the different treatments investigated in this study: (A-B) reference beer (low ethanol level: 5.5 v/v% and low iso-α-acid level: 13 ppm and 17 IBU), (C-D) beer with medium ethanol level (8 v/v%), (E-F) beer with high ethanol level (11 v/v%), (G-H) beer with medium iso-α-acid level (35 ppm and 50 IBU), and (I-J) beer with high iso-α-acid level (170 ppm and 135 IBU). Data are presented as the average of three biological replicates (*n* = 3) and error bars represent the standard error of the mean.

**Figure S2**: Heatmap visualizing the ratio between the concentration of the different chemical compounds in the negative controls (without microbial community) at day 60 and the concentrations measured at the start (day 0) of the maturation, indicating whether compound concentrations in the negative controls increased (red) or decreased (blue) throughout wood maturation. Results are presented for the five different treatments (*n* = 3), including: reference beer (low ethanol level: 5.5 v/v% and low iso-α-acid level: 13 ppm and 17 IBU), beer with medium ethanol level (8 v/v%), beer with high ethanol level (11 v/v%), beer with medium iso-α-acid level (35 ppm and 50 IBU), and beer with high iso-α-acid level (170 ppm and 135 IBU). Welch t-tests were performed to test whether the changes in concentration throughout wood maturation of the negative controls were significant. *P*-values obtained in the t-tests are presented in the cells: *\* p* < 0.05, ** *p* < 0.01, *** *p* < 0.001.

**Figure S3**: Temporal dynamics in microbial community composition assessed via cultivation on different media. For the composition of the media as well as the incubation conditions, the reader is referred to Table S9 (Supplementary Information). Results are presented per treatment: (A) ethanol treatments (low level: 5.2 v/v%, medium level: 8 v/v%, high level: 11 v/v%) and (B) iso-α-acid treatments (low level: 13 ppm and 17 IBU, medium level: 35 ppm and 50 IBU, high level: 170 ppm and 135 IBU).

## 9 Data Availability Statement

The data used in this study (qPCR and chemical data) are available in Supplementary Information. Sequences of the 16S rRNA gene (bacteria), and the 28S rRNA gene or the ITS1-5.8S rRNA-ITS2 region (fungi) of the microbial strains investigated in this study have been deposited in GenBank under the following accession numbers: OM432146 (*A. malorum*); OM432147 (*G. oxydans*); OM432145 (*L. brevis*); OM432144 (*P. damnosus*); OM432156 (*S. cerevisiae*); OM432150 (*B. bruxellensis*); OM432151 (*P. membranifaciens*); and OM432152 (*C. friedrichii*).

