## Supplementary figures and images for "Beer ethanol and iso-α-acid level affect microbial community establishment and beer chemistry throughout wood maturation of beer"

### Fig S1 Line plots per treatment

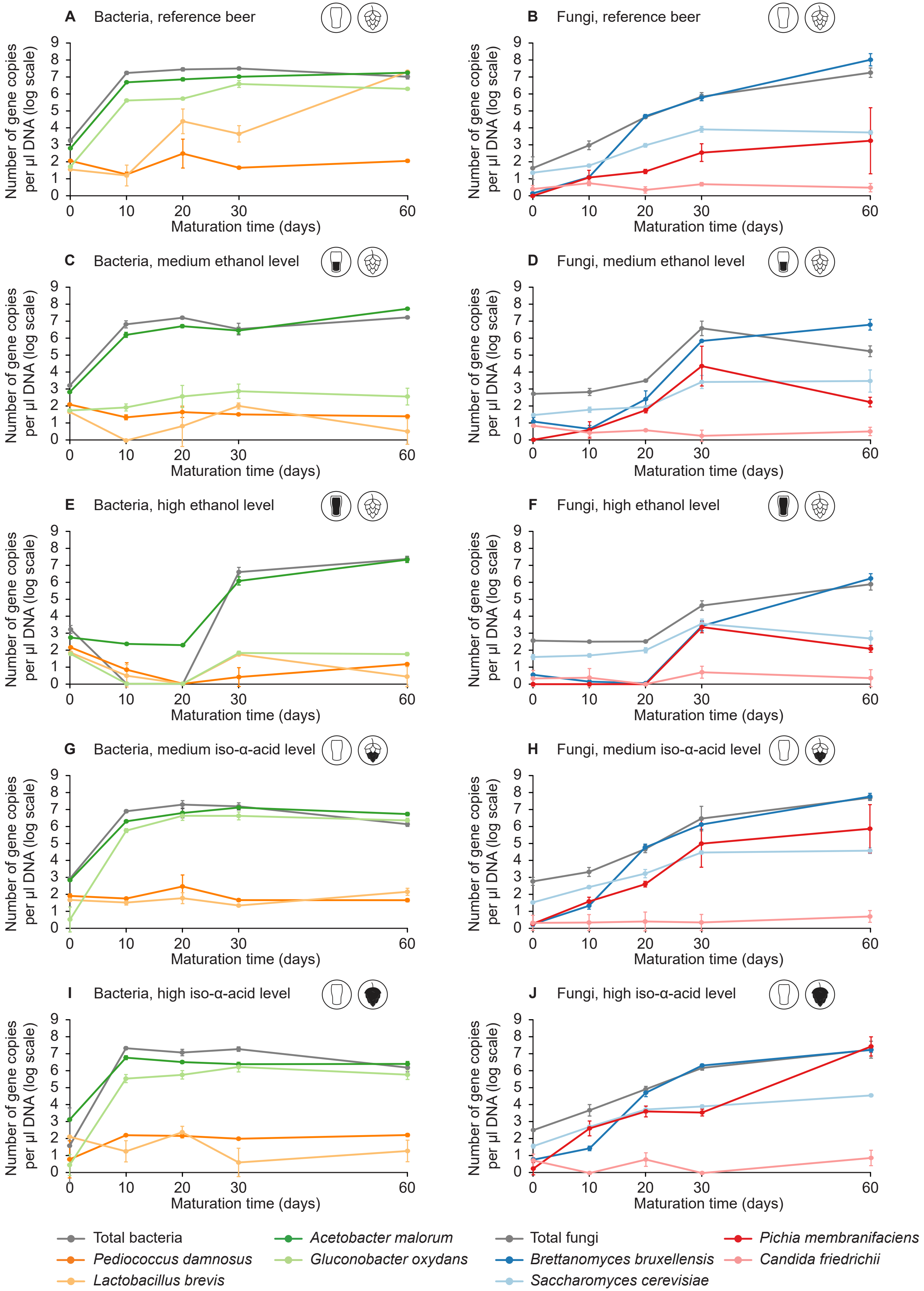

### Fig S2 Chemistry heatmaps negative controls

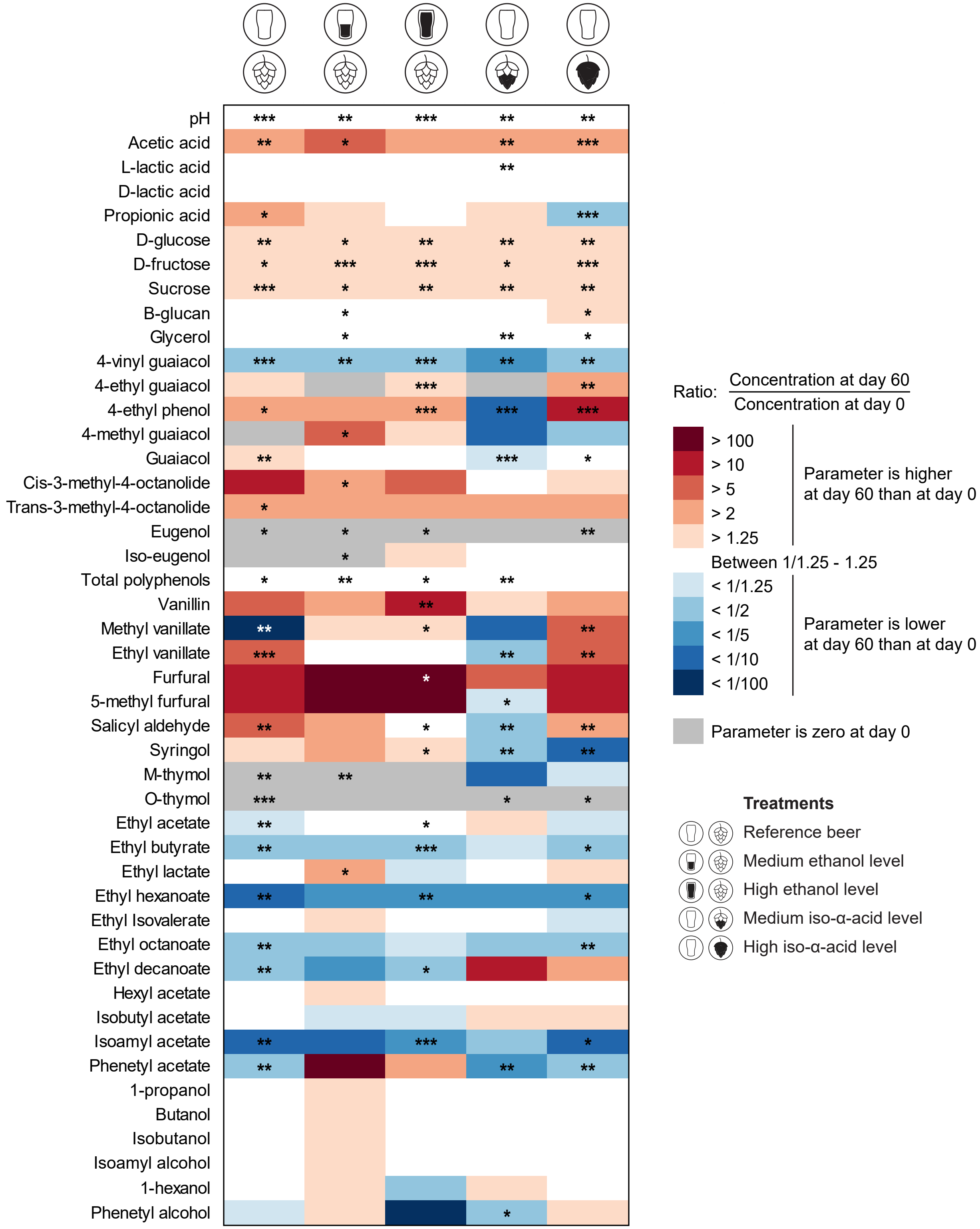

### Fig S3 Plate counts

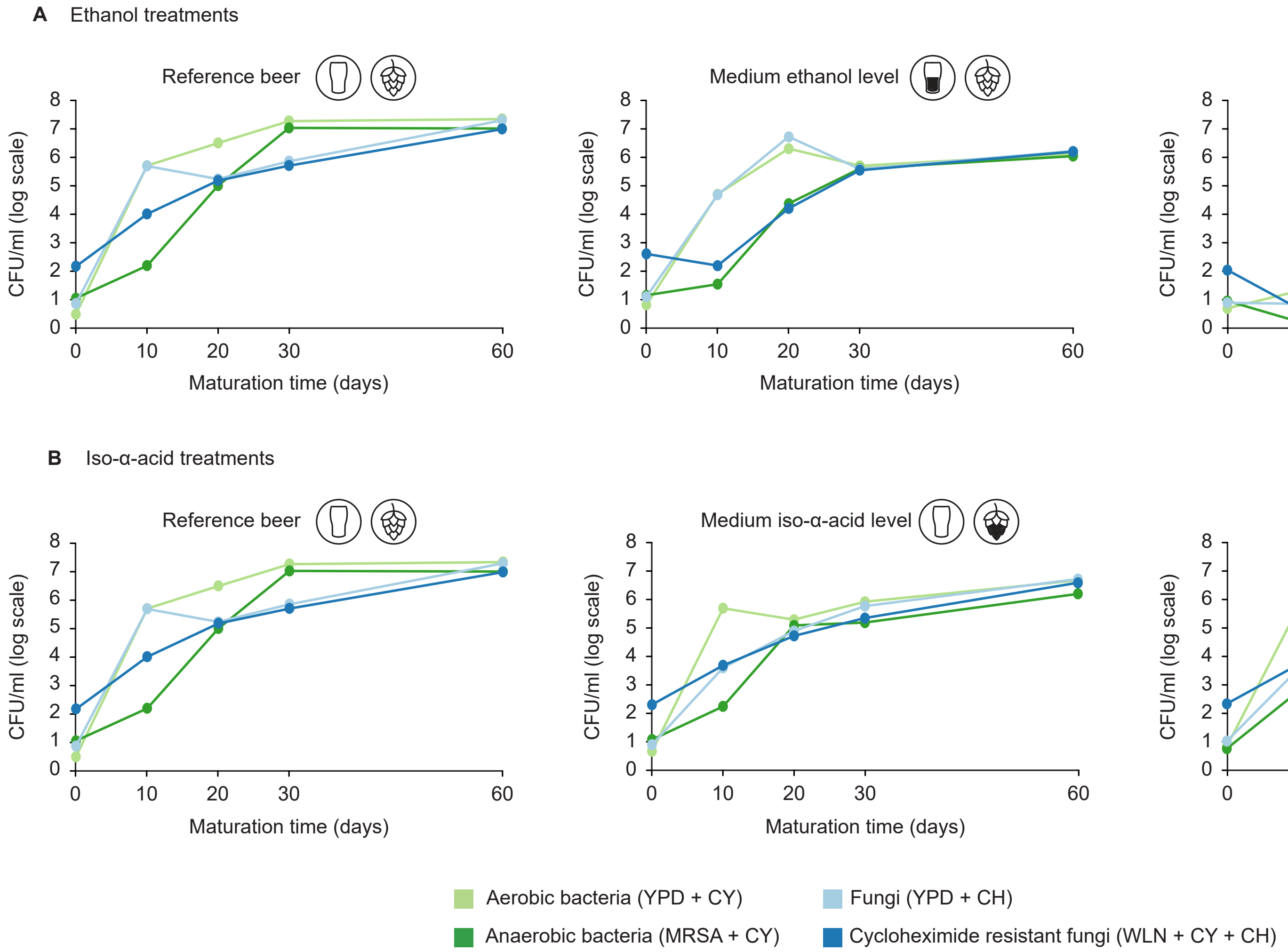
